# The revised reference genome of the leopard gecko (*Eublepharis macularius*) provides insight into the considerations of genome phasing and assembly

**DOI:** 10.1101/2023.01.20.523807

**Authors:** Brendan J. Pinto, Tony Gamble, Chase H. Smith, Shannon E. Keating, Justin C. Havird, Ylenia Chiari

**Affiliations:** School of Life Sciences, Arizona State University, Tempe, AZ USA; Center for Evolution and Medicine, Arizona State University, Tempe, AZ USA; Department of Zoology, Milwaukee Public Museum, Milwaukee, WI USA; Department of Biological Sciences, Marquette University, Milwaukee WI USA; Bell Museum of Natural History, University of Minnesota, St Paul, MN USA; Department of Integrative Biology, University of Texas at Austin, Austin, TX, USA; Department of Biology, George Mason University, Fairfax, VA, USA

## Abstract

Genomic resources across squamate reptiles (lizards and snakes) have lagged behind other vertebrate systems and high-quality reference genomes remain scarce. Of the 23 chromosome-scale reference genomes across the order, only 12 of the ~60 squamate families are represented. Within geckos (infraorder Gekkota), a species-rich clade of lizards, chromosome-level genomes are exceptionally sparse representing only two of the seven extant families. Using the latest advances in genome sequencing and assembly methods, we generated one of the highest quality squamate genomes to date for the leopard gecko, *Eublepharis macularius* (Eublepharidae). We compared this assembly to the previous, short-read only, *E. macularius* reference genome published in 2016 and examined potential factors within the assembly influencing contiguity of genome assemblies using PacBio HiFi data. Briefly, the read N50 of the PacBio HiFi reads generated for this study was equal to the contig N50 of the previous *E. macularius* reference genome at 20.4 kilobases. The HiFi reads were assembled into a total of 132 contigs, which was further scaffolded using HiC data into 75 total sequences representing all 19 chromosomes. We identified that 9 of the 19 chromosomes were assembled as single contigs, while the other 10 chromosomes were each scaffolded together from two or more contigs. We qualitatively identified that percent repeat content within a chromosome broadly affects its assembly contiguity prior to scaffolding. This genome assembly signifies a new age for squamate genomics where high-quality reference genomes rivaling some of the best vertebrate genome assemblies can be generated for a fraction previous cost estimates. This new *E. macularius* reference assembly is available on NCBI at JAOPLA010000000. The genome version and its associated annotations are also available via this Figshare repository https://doi.org/10.6084/m9.figshare.20069273.

## Introduction

Genomic data in squamate reptiles (lizards and snakes) has lagged behind other vertebrate model systems, such as birds and mammals, and high-quality reference genomes remain scarce (Bravo et al. 2022; Hotaling et al. 2021; Pinto et al. 2023). Of the 23 previously published chromosome-scale squamate reference genomes, only 12 of the ~60 families are represented. Within geckos, chromosome-level genomes are exceptionally sparse, representing only two of the seven extant gecko families (Pinto et al. 2022; Yamaguchi et al. 2021). While there are also a handful of non-chromosome level gecko genomes, including the leopard gecko, *Eublepharis macularius* (Gekkota: Eublepharidae), draft genomes are limited in their utility to address many ecological and evolutionary hypotheses (Xiong et al. 2016). Adding to the current genomics resources in geckos, we used the latest advances in genome sequencing and assembly methods to generate one of the highest quality squamate reference genomes to date for *E. macularius*.

Investigating the evolution of genomes and phenotypes involves examining multiple species in a phylogenetic context. The foundation of integrative and comparative biology is that one can infer the likely ancestral condition for a specific trait, such as gene structure or function, by carefully choosing species that span the deepest bifurcations of a particular clade of interest (Felsenstein, 1985; Bryant and Russell, 1992; Witmer, 1995; Pagel et al. 2004). This idea is also justification of using model species (Wake, 2008; Hall, 2012; Dobzhansky, 1973; Sanger and Rajakumar, 2019). Consequently, gecko lizards (Infraorder Gekkota) are prime candidates to be a powerful model system for vertebrate genomics. Geckos are a species rich group of lizards— 2,186 species as of December 2022—distributed in tropical and subtropical regions around the world (Uetz et al. 2021; Bauer, 2013). Geckos make up a large part of amniote diversity, representing ~8% of total species. They are the sister clade to all other lizards and snakes, with the possible exception of the poorly known, limbless, dibamids—whose phylogenetic position remains unresolved (Wiens et al. 2012; Zheng and Wiens, 2016; Townsend et al. 2004). Indeed, geckos diverged from all other lizards and snakes over 250 million years ago and extant geckos began to diversify ~120 million years ago (Gamble et al. 2011 & 2015a). For scale, this makes geckos as divergent from other squamates as humans are from a platypus (Kumar et al. 2017). Therefore, the inclusion of geckos in any evolutionary study of squamates is crucial to understanding genome evolution in lizards and snakes more broadly. As such, a high-quality genome assembly from a gecko is an important resource for investigating amniote genome evolution. Geckos are also interesting in their own right, for example, geckos possess unique biological traits, many of which have evolved repeatedly within the group, including adhesive toepads, sex determination systems, and photic activity patterns (Gamble et al. 2012; 2015a; 2015b; 2019; Pinto et al. 2019); and deep investigations into these, and other, aspects of gecko biology requires robust genomic resources.

Since its humble beginnings as a charismatic staple in the international pet trade, *Eublepharis macularius* has become a standard laboratory model system for studying a variety of biological questions surrounding tissue regeneration, coloration, sex determination, behavior, and cancer (Whimster, 1965; Viets et al., 1993; McLean & Vickaryous, 2011; Sakata et al., 2022; Delorme et al., 2012; Kiskowski et al., 2019; Szydłowski et al. 2020; Glimm et al. 2021; Guo et al. 2021; Agarwal et al. 2022; Katlein et al. 2022). However, more detailed investigations into genotype-phenotype associations in *E. macularius* have been hampered by modest genomic resources (Chernyavskaya et al. 2022; Gamble, 2019; Nurk et al. 2022). Thus, available genomic resources remain a research limitation and a high-quality reference genome for this model taxon will reduce potential error in downstream inference (Kim et al. 2019). Here, we generated a phased, chromosome-level genome assembly using a combination of Pacific Biosciences® High Fidelity (PacBio HiFi) and Dovetail® Omni-C (HiC) data. This assembly stands as one of the best primary assemblies for any squamate (132 contigs), and perhaps any vertebrate, standing alongside the highest-quality assemblies like the newest Telomere-to-Telomere assembly for humans (Nurk et al. 2022).

## Methods

### Data Generation

The reference genome individual was an unsexed, juvenile *E. macularius* (TG4126) incubated at a female temperature. This individual was homozygous for the recessive Tremper strain albino allele and heterozygous for the incomplete dominant Lemon Frost allele. We also sequenced the parents for phasing using the trios approach: the sire (TG4151) was homozygous for the recessive Tremper strain albino allele, heterozygous for the recessive Murphy patternless allele, and a Tremper giant - a phenotype with unknown genetics; the dam (TG4152) was homozygous for the recessive Tremper strain albino allele, heterozygous for the recessive Murphy patternless allele, and heterozygous for the incomplete dominant Lemon Frost allele (Supplemental Figure 1). We extracted high molecular weight DNA from blood of the offspring by Salting Out Phenol-Chloroform with an Ethanol Precipitation (SOP-CEPC; Pinto et al. 2021) and sent DNA to the Genomics & Cell Characterization Core Facility (GC3F) at the University of Oregon. A single PacBio HiFi library was sequenced across 3 SMRT cells. We generated a HiC library from the same individual using a DoveTail® Omni-C kit (Cantata Bio; Cambridge, MA, USA) and sequenced on an Illumina NovaSeq 6000 at the Texas A&M Agrilife Core Facility (College Station, TX, USA). DNA from the sire and dam were extracted using the Qiagen® DNeasy Blood and Tissue kit and sequenced on an Illumina NovaSeq 60000 by Novogene® (Beijing, China). All Illumina® sequencing was run using PE 150 bp (2×150) reads.

### Genome Assembly

We generated five total assemblies from using offspring HiFi data using HiFiasm [v0.16.1-r375] (Cheng et al. 2021 & 2022). The assemblies were as follows (1) an aggregate contig assembly using all HiFi reads (this assembly was chosen for further scaffolding of the reference genome), (2-3) the trio phased assemblies (paternal and maternal) using parental short reads, (4-5) the HiC phased assemblies (haplotypes 1 and 2). Although there were subtle differences in contiguity between primary haplotype assemblies, they were overall very similar in quality (Table 1). After the initial contig assemblies, we scaffolded contig set (1) using the offspring HiC data in 3D-DNA [v201008] (Dudchenko et al. 2017). We visualized the final HiC contact map for misassemblies and, with no large-scale misassemblies visible, we manually refined the contact map using Juicebox Assembly Tools [v1.11] (Durand et al. 2016).

**Table 1:**
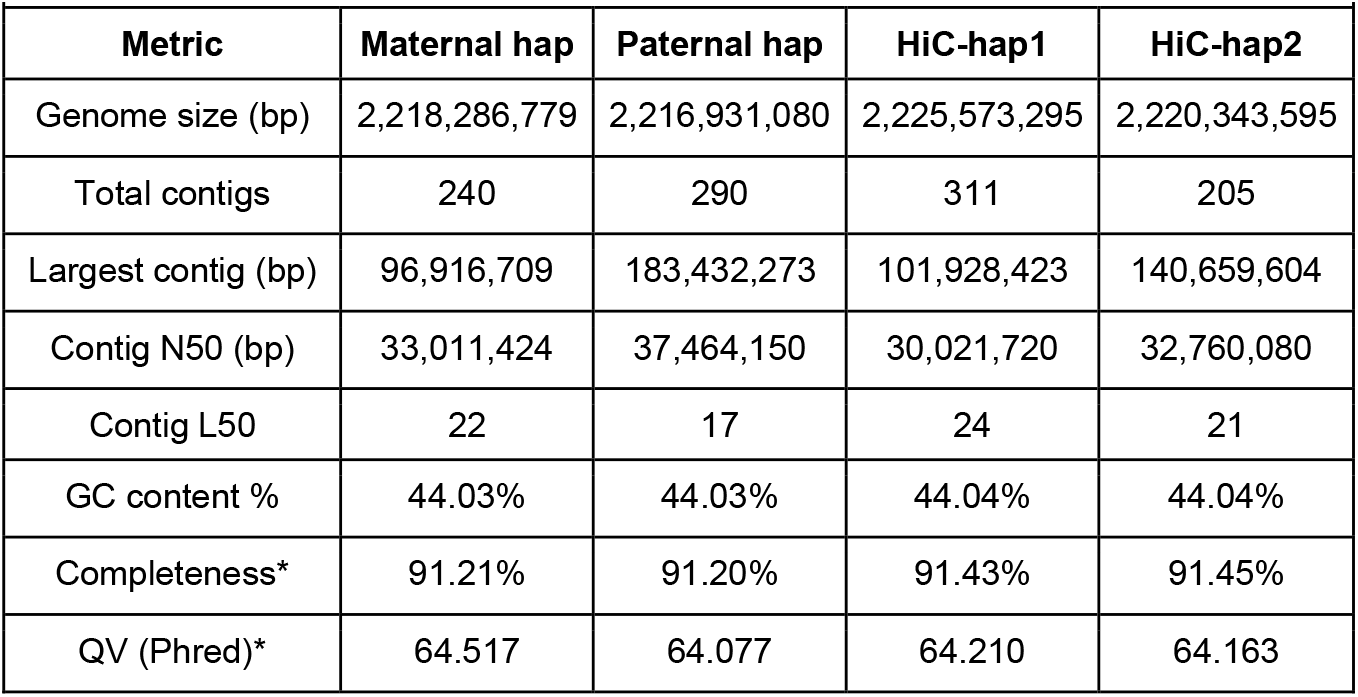
Assembly metrics comparing previous different phasing schemes for the assembled HiFi reads. * = merqury scores

### Genome QC

We estimated metrics of genomic completeness using the raw sequencing reads and a database of conserved single-copy orthologs with merqury [v1.3.0] (Rhie et al. 2020) and Benchmarking Universal Single-Copy Orthologs (BUSCO) [v5.1.2] (Simão et al. 2015), respectively. We implemented all BUSCO analyses using the gVolante web server [v2.0.0] (Nishimura et al. 2017) with the Core Vertebrate Genes (CVG) and Sauropsida_odb10 databases. We also compared the relative ability of the two methods of phasing available in Hifiasm, parental data (trios) or chromatin-contact data (HiC) using merqury. We counted kmers for the offspring HiFi data, as well as the parental Illumina data using meryl [v1.3]. To calculate genomic heterozygosity of parental samples, we mapped reads to the reference using bwa-mem2 [v2.2.1] (Vasimuddin et al. 2019) and called SNPs using freebayes [v1.3.5] (Garrison and Marth, 2012). We removed non-biallelic sites, sites with <30 quality score, and sites with a read depth <5 using vcftools [0.1.14-12] (Danecek et al. 2011).

### Genome Annotation

We masked the assembly for repeats using a combination of RepeatModeler [v2.0.3] and RepeatMasker [v4.1.2] (Flynn et al. 2020; Smit et al. 2013). Later statistical analysis of the genomic repeat content used Mann-Whitney-Wilcoxon tests (Figure 2). For the initial release of the genome assembly, we chose to liftover the annotations from the previous reference genome (Xiong et al. 2016), given the absence of additional RNAseq data for *Eublepharis macularius.* Previous annotations were transferred using Liftoff [v1.6.3] (Li, 2018; Shumate & Salzberg, 2020). We diagnosed success by comparing the transferred annotations to the original annotation using BUSCO (Table 2).

**Table 2:** Assembly metrics comparing previous *Eublepharis macularius* reference genome (Xiong et al. 2016) to MPM_Emac_v1.0 reference assembly. BUSCO scores abbreviated as follows, C=complete, S=complete single copy, D=complete multi-copy, F=fragmented, M=missing. * missing data not included in calculation

| Metric | Xiong et al. 2016 | MPM_Emac_v.1.0 |
| --- | --- | --- |
| Contig N50 (bp)* | 20,426 | 80,105,973 |
| Contig L50* | 27,671 | 9 |
| Scaffold N50 (bp) | 663,762 | 145,573,841 |
| Scaffold L50 | 796 | 6 |
| Repeat content % | 42.1% | 48.0% |
| GC content % | 43.6% | 44.1% |
| Total scaffolds | 206,349 | 75 |
| Genome Size (bp)* | 1,981,651,937 | 2,237,374,058 |
| BUSCO Sauropsida (Genome) | C:93.4%[S:91.4%,D:2.0%], F:2.0%,M:4.6%,n:7480 | C:95.1%[S:93.0%,D:2.1%],F:0.9%,M:4.0%,n:7480 |
| BUSCO Sauropsida (Annotation) | C:90.6%[S:88.7%,D:1.9%], F:3.2%,M:6.2%,n:7480 | C:89.3%[S:87.6%,D:1.7%],F:3.6%,M:7.1%,n:7480 |
| BUSCO CVG (Genome) | C:96.6%[S:95.7%,D:0.9%], F:2.1%,M:1.3%,n:233 | C:99.2%[S:97.9%,D:1.3%],F:0.9%,M:0.1%,n:233 |

## Results and Discussion

The HMW DNA extraction optimized specifically for extremely small sample inputs (Pinto et al. 2021 & 2022), e.g. *Sphaerodactylus* gecko tissues, worked well in our *E. macularius* tissue samples. DNA extractions had an average molecule length of ~52 kilobase-pairs (kb) and 45% of the total extraction >50kb, much longer than the input for PacBio HiFi DNA sequencing allowed for pre-library preparation shearing optimization (Supplemental Figure 2). Post-circular consensus sequencing (CCS) correction, we recovered 66Gb of data (~30X coverage) with an average read length of 19.6kb and a read N50 of 20.4kb. With a read length N50 equal to the contig N50 of the previous *E. macularius* reference genome, the primary assembly generated by HiFiasm contained 132 total contigs with a N50 and L50 of 80,105,973bp and 9, respectively. The smallest contig was 2,547bp and the largest was 188,850,821bp. Scaffolding using HiC data added 61 gaps to produce the final assembly. The final *E. macularius* genome assembly contained 19 primary chromosome-level scaffolds and 56 unanchored contigs, ranging from ~400kb to ~11kb (75 total sequences), with a scaffold N50 and L50 of 145,573,841bp and 6, respectively.

RepeatModeler identified 47.98% of the genome as repetitive (Table 2). Most repetitive elements in the genome remain unclassified (21.1%), followed by a majority being retroelements, either LINEs (14.31%), LTR elements (4.96%), or SINEs (4.06%). All other categories combined totaled <4% of the total repetitive elements, including DNA transposons (1.91%). We calculated the merqury completeness score at 91.5% using the PacBio HiFi reads used to generate the primary assembly, suggesting our assembly was largely complete. The BUSCO completeness scores were comparably valued at 99.2% and 95.1% using the Core Vertebrate Genome (CVG) and Sauropsida ortholog databases, respectively.

Nine of the 19 chromosomes were assembled as single contigs (Figure 1). With such high contiguity of the primary assembly prior to scaffolding, we further examined which chromosomes we assembled as a single contig. Centromeres are a typical assembly breakpoint (Peona et al. 2020) in genome assemblies but *E. macularius* has a karyotype consisting of 19 pairs of acrocentric chromosomes gradually decreasing in size (Gorman, 1973). Like other geckos, *E. macularius* chromosomes possess no sharp divide between macro- and micro-chromosomes (Figure 1; Pinto et al. 2023). Thus, greater contiguity may be due, in part, to this chromosomal arrangement. We aimed to identify any properties of individual chromosomes that might explain the contiguity of the assembled molecules. We binned chromosome-length scaffolds into single-contig and multi-contig categories and compared these groups with relation to their GC content and repetitive DNA content (Figure 2). We chose to use these statistics considering GC content is often considered a proxy for DNA stability and repetitive elements have been consistently demonstrated to cause assembly gaps (Eyre-Walker and Hurst, 2001; Peona et al. 2020). Interestingly, we did not find any significant differences between the two groups for either comparison. We did observe a higher repetitive element content in multi-contig scaffolds, as we expected, but the difference was not found to be significant (p = 0.14). However, the lack of significance could easily be explained by the small sample size (i.e. N=19, the number of chromosomes present) and qualitatively there may be a lower repeat content in those chromosomes assembled as a single contig than those scaffolded together as multiple contigs (Figure 2). Alternatively, these (rare) assembly gaps may be caused by additional genomic elements that are beyond the scope of the present study.

**Figure 1:**
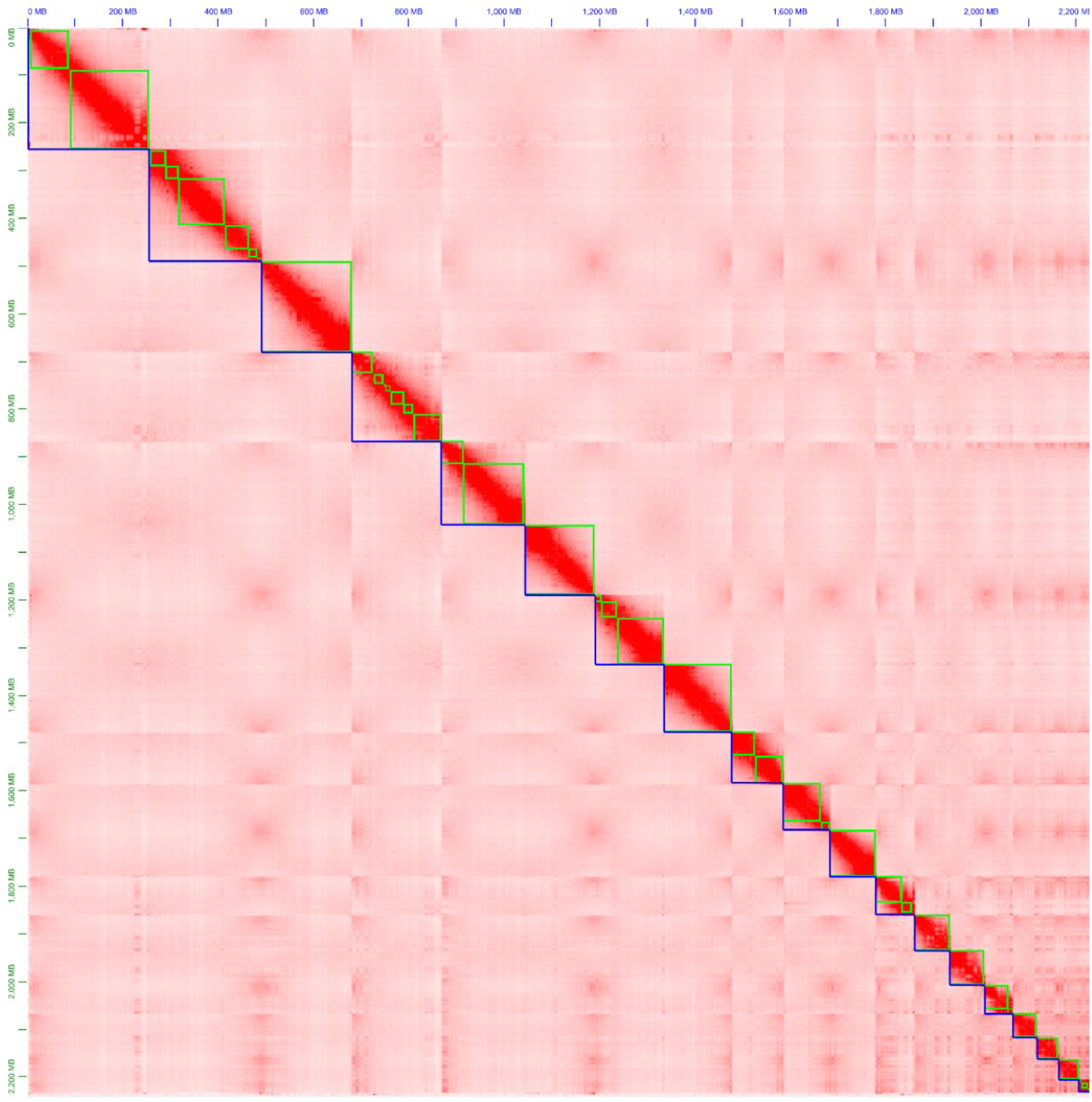
HiC contact map for the MPM_Emac_v1.0 assembly. Each blue segment indicates the delimitation of a chromosome-length scaffold, while the internal green squares indicate contigs. Approximately half of the assembled chromosomes are represented by a single contig, indicating the extreme contiguity of the primary assembly pre-scaffolding.

**Figure 2:**
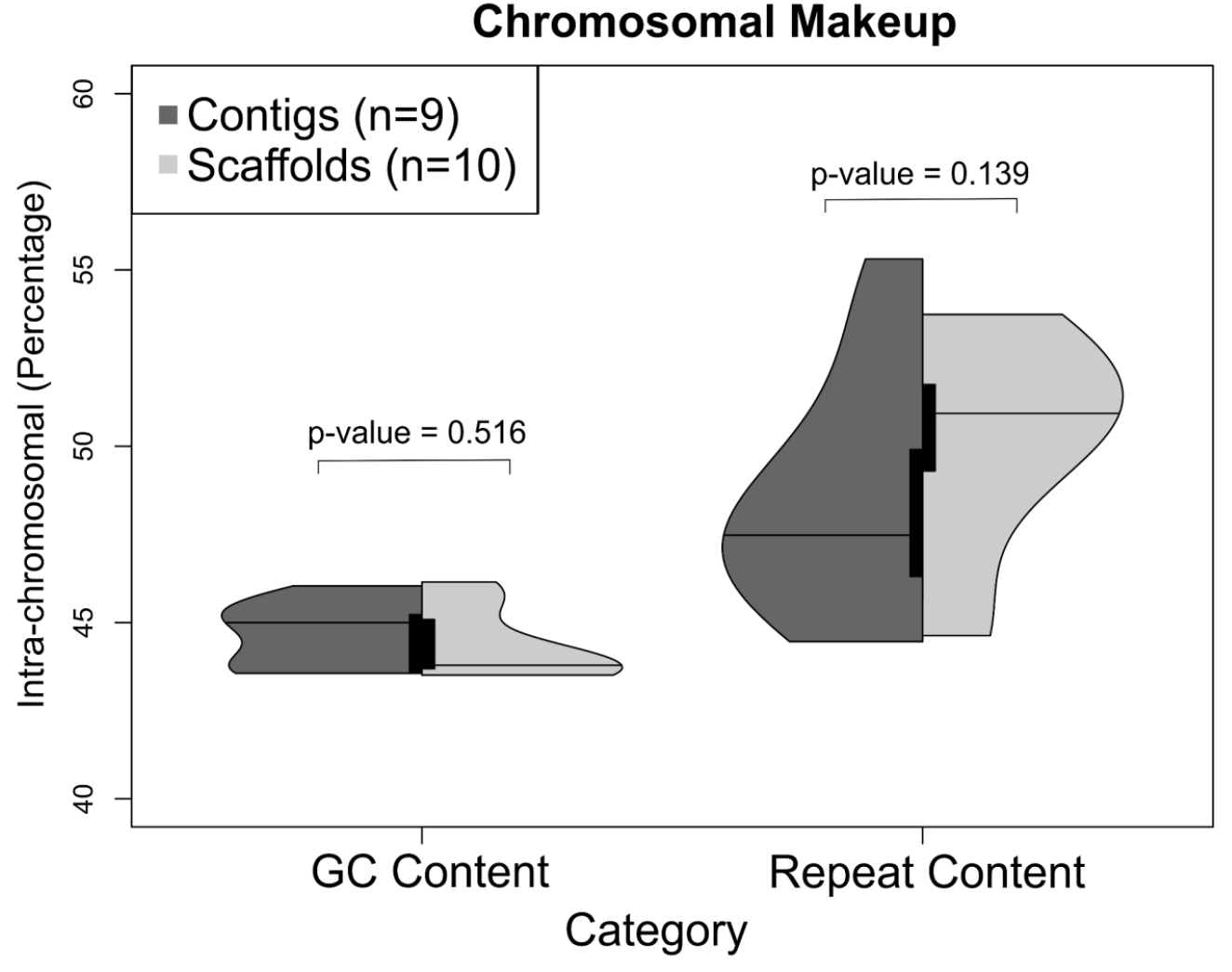
Comparison between chromosomes assembled as a single contig (“Contigs”, dark gray) and those composed of multiple contigs (“Scaffolds”, light gray) displayed using vioplot (Adler et al. 2022). The violins represent the distribution of the underlying data points, while the internal bars represent a traditional bar graph representation. The mid-lines represent the median of the data. Our *a priori* hypothesis was that chromosomes assembled as a single contig would possess lower overall GC content and/or repeat content. However, neither GC content or repetitive element content were significantly different using Mann-Whitney-Wilcoxon tests. Qualitatively, there appears to be a difference in median repeat content between the two groups. It’s possible that our ability to detect a true difference using frequentist methods lies in the low sample size (N=19).

There is an inherent tradeoff to account for when planning a genome assembly and phasing experiment. Indeed, low heterozygosity tends to improve contiguity of the final assembly, but heterozygosity is a necessary component to successful phasing (e.g. Chin et al., 2016; Koren et al. 2018). We investigated the phasing capabilities of the parent/offspring trio approach to a single-individual with HiC data in *E. macularius,* an animal with low overall heterozygosity and no sex chromosomes. Perhaps surprisingly, HiC outperformed the Trios method for phasing, where phase blocks are approximately equal to contig sizes (Figure 3A5-6, B-5-6). However, neither method provided complete haplotype-resolution with either (1) high switch error rates disrupting contig phasing (Trios, Figure A3-4) or (2) inconsistent assignment to maternal/paternal haplotypes (HiC, Figure B3-4).

**Figure 3:**
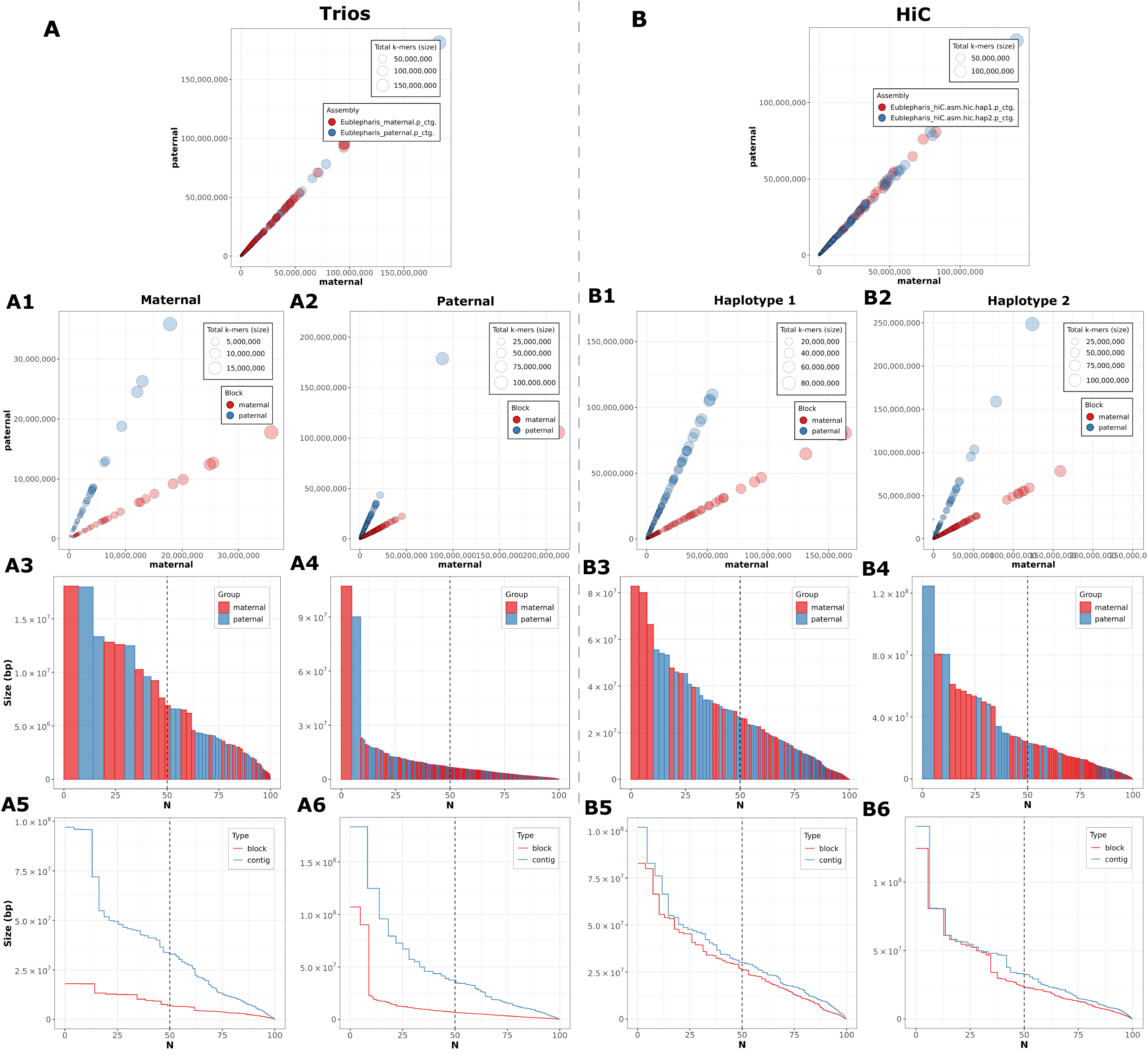
Comparative QC results from assembly phasing generated by merqury (Rhie et al. 2020) between (A) Trios phasing and (B) HiC phasing methods. HiC equaled or outperformed Trios phasing in all measured categories. Notably, Trios phasing appears to have suffered from high switch-error rates, which resulted in short phase block, relative to contig size. HiC phasing performed extremely well, however by definition; HiC phasing was unable to coordinate multiple phased contigs to their parent of origin.

We hypothesized that HiC outperformed the trios approach because of low levels of heterozygosity contained within this lab bred lineage of leopard gecko—originally sourced from the pet trade. To examine this further, we mapped reads from each parent to the reference, called SNPs (see methods), and estimated heterozygosity by dividing the total number of SNPs by the total genome size (father/TG4151 = 0.35%; mother/TG4152 = 0.38%). However, we acknowledge that only heterozygous sites that are not shared between parents are informative for phasing purposes. We identified sites that were not shared between parents using vcf-compare to calculate the informative heterozygosity rate for phasing (father/TG4151 = 0.15%; mother/TG4152 = 0.18%). Indeed, less than 50% of heterozygous sites in each parent are informative for phasing, which limits theoretically informative phasing sites to <2,500 SNPs per Mb on average. This constraint on informative sites was not observed in HiC phasing given that every heterozygous site in the genome is theoretically informative for HiC phasing – approximately doubling the number of informative sites when phasing with HiC data. In sum, this genome assembly experiment was conducted on a trio of animals with too little heterozygosity for successful offspring phasing, but HiC provided sufficient resolution for phasing. For future studies facing a similar situation, we suggest either planning the experiment around a single individual using HiC or outcrossing two individuals with different genetic backgrounds and sequencing this doubly heterozygous offspring trio to increase site informativeness, which are analogous to the established standards for traditional linkage mapping experiments (e.g. Amores et al. 2014).

Our annotation for this reference genome, MPM_Emac_v1.0, maintained a completeness of 89.3% using the sauropsida_odb10 dataset in BUSCO [v5.1.2] (Simão et al. 2015), nearly mirroring the Xiong et al. (2016) original reference genome annotation of 90.6%. Of note, these numbers do not match those from the Xiong et al. (2016) manuscript due to changes in both software versions and query databases. We also compared other differences between the current assembly and the original reference assembly. MPM_Emac_v1.0 assembly size ~12% larger than Xiong et al. (2016)—2.24Gb vs. 2.02Gb, respectively. Interestingly, MPM_Emac_v1.0 is much closer to the kmer estimated genome size from Xiong et al. (2016) of 2.23Gb. There is also an increase of repetitive DNA content in MPM_Emac_v1.0 of ~6% (Table 2). However, the GC content deviated by 0.5% between the two assemblies, indicating that the GC content in gecko genomes may not be as biased with short-read based sequence data as might be anticipated *a priori* (Benjamini and Speed, 2012).

In conclusion, we present a chromosome-level genome assembly for the leopard gecko, *E. macularius.* This is simultaneously the first phased chromosome-level assembly and the first long-read based genome assembly available for any species of gecko. Further, this assembly is one of the most contiguous squamate genomes available and has achieved the second highest BUSCO score of any squamate genome (Pinto et al. 2023). The last hurdle for this assembly to overcome before this assembly can be considered a finished “telomere-to-telomere” assembly is placing the final 5.02Mb of unassembled sequence into the 19 primary scaffolds representing the 19 chromosomes of *E. macularius.* This would likely require generation of a modest number of ultra-long reads to fill gaps and complete centromeric/telomeric regions (e.g. Rautiainen et al. 2022). Nonetheless, our genome assembly represents the new ‘gold standard’ in squamate genomes at this ever-fleeting moment.

## Data Availability

The raw data generated in this study has been deposited to NCBI SRA under Bioproject PRJNA884264. The genome version described in this study MPM_Emac_v1.0,annotation file, and all four phased assemblies are available via this Figshare repository https://doi.org/10.6084/m9.figshare.20069273. Additionally, genome assembly version MPM_Emac_v1.0.1 is archived as a Whole Genome Shotgun project deposited at DDBJ/ENA/GenBank under the accession JAOPLA000000000. The genome version and its associated annotations are also available via this Figshare repository https://doi.org/10.6084/m9.figshare.20069273.

## Acknowledgements

The authors would like to thank: M. Weitzman, formally at the University of Oregon GC3F sequencing core, for her exceptional help and support of the HiFi sequencing portion of this project; R. Tremper for generously providing the parental geckos; and Gamble Lab animal care technicians. Gecko breeding and tissue sampling were conducted under Marquette University IACUC protocol AR-279.

## Supplemental Information

**Supplemental Figure 1:**
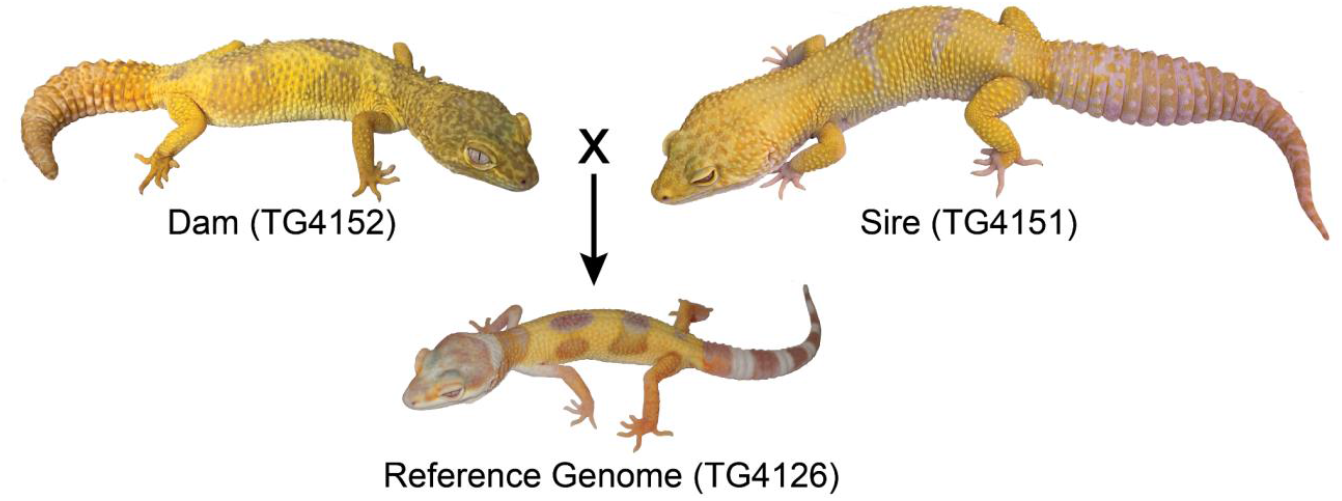
Crossing scheme to generate the reference genome animal for *Eublepharis macularius,* TG4126.

**Supplemental Figure 2:**
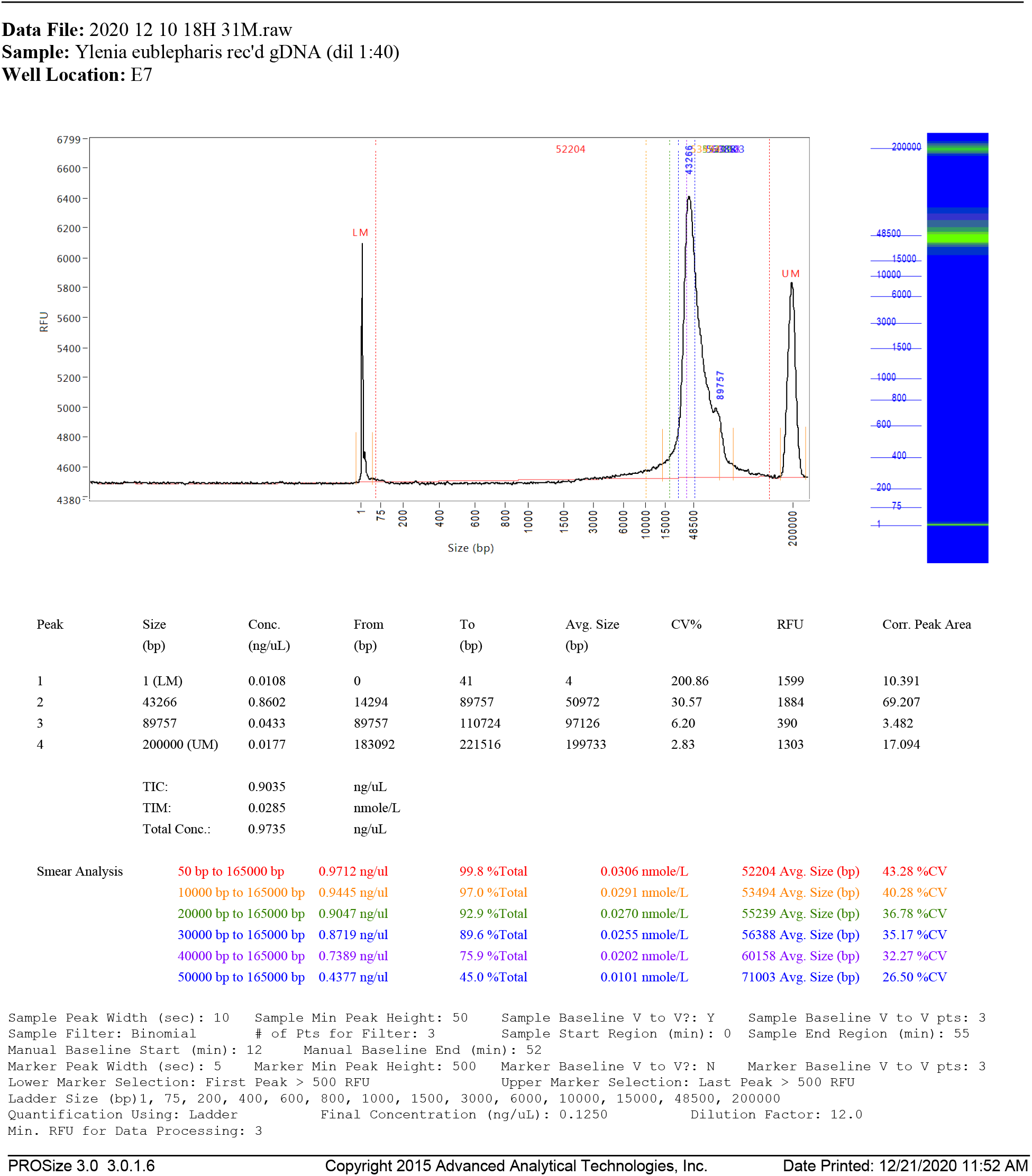
High-molecular weight DNA extraction Bioanalyzer results.

